# *Cnes2b* Regulates Host Resistance, Inflammatory Responses and Tissue Damage Following *Cryptococcus deneoformans* Infection

**DOI:** 10.1101/2024.09.14.613072

**Authors:** Isabelle Angers, Annie Beauchamp, Marwa El Sheikh, Eva Kaufmann, Donald C. Vinh, Salman T. Qureshi

## Abstract

The 32.1 Mb *Cnes2* chromosome 17 interval was shown to confer resistance to progressive *Cryptococcus deneoformans* 52D infection. To refine the location of *Cnes2* host resistance genes, a subcongenic mouse strain (B6.CBA-*Cnes2b*) that contains 8.7 Mb from the telomeric region of *Cnes2* was created. At 28 days postinfection B6.CBA-*Cnes2b* mice had a lower lung fungal burden, increased lung injury, as well as mortality compared to C57BL/6N. B6.CBA-*Cnes2b* mice had increased pulmonary production of pro-inflammatory mediators, chemokines and Th1-type cytokines as well as increased recruitment of monocytes and neutrophils to the lungs. *Cnes2b* also regulated several elements of the host response to *C. deneoformans* 52D infection in a sex-dependent manner. Specifically, male B6.CBA-*Cnes2b* mice had a lower lung fungal burden, increased brain injury and mortality relative to females. Taken together these findings demonstrate that *Cnes2b* regulates host inflammation in a manner that controls fungal burden and increases tissue damage. Precise identification of the genes encoded by *Cnes2b* could reveal key mechanisms of cryptococcal host resistance and immune reconstitution or postinfectious inflammatory syndromes.

**Importance:** The 32.1 Mb *Cnes2* congenic interval from chromosome 17 of resistant CBA/J mice regulates host resistance to *C. deneoformans* 52D infection. This study characterizes the host response of B6.CBA-*Cnes2b* mice that carry an 8.7 Mb sub-congenic interval derived from *Cnes2* following *C. deneoformans* 52D infection. B6.CBA-*Cnes2b* mice had reduced lung fungal burden, increased lung and brain injury, and mortality. The effects of *Cnes2b* differed between male and female subcongenic mice and are consistent with known sex differences in human cryptococcal disease. The host response of B6.CBA-*Cnes2b* mice reflects a crucial balance between effective control of fungal burden and potentially deleterious consequences of enhanced inflammation during cryptococcal infection as predicted by the damage response framework. Further analysis of the *Cnes2b* sub-congenic interval will lead to definitive identification of genes that confer resistance to progressive cryptococcal infection and/or contribute to deleterious inflammatory responses. Defining key mechanisms that regulate the immune response to *Cryptococcus sp.* is an important step towards the development of host-directed therapeutics that could improve disease outcomes.

## Introduction

*Cryptococcus deneoformans* is a soil-associated encapsulated yeast that causes human infection via the respiratory tract (1). Frequently, cryptococcal lung infection may be asymptomatic or cause mild disease and may resolve without specific therapeutic intervention (2). Conversely, among severely immunocompromised patients, *Cryptococcus sp.* frequently disseminates from the lungs to the central nervous system and causes potentially fatal meningitis (3). In areas with high rates of HIV infection *Cryptococcus sp.* are the most common cause of adult meningitis, accounting for 13-24% of AIDS-associated deaths and resulting in significant long-term morbidity among many survivors (4). In resource rich areas with access to highly effective antiretroviral therapy, one-third of deaths occur in non-HIV-infected patients (5, 6). Risk factors for the severe cryptococcal disease among non-HIV-infected patients include liver or kidney disease, transplantation, sarcoidosis, cellular or humoral immune defects, and the use of corticosteroids or novel immunomodulatory therapies (6). Natural susceptibility to human cryptococcal disease has also been associated with rare inborn errors of immunity (4). Finally, genetic polymorphisms of Fc-gamma receptors, mannose-binding lectin, Dectin-2, Toll-like receptors and macrophage colony-stimulating factor have been associated with cryptococcal infection, although the mechanisms by which these variants influence disease progression remain elusive (7).

The outcome of cryptococcal infection is determined by a complex interplay between fungal virulence characteristics and host immune responses. Cryptococcal disease is a prototypical example of the Damage-Response Framework (DRF) of microbial pathogenesis (8). According to the DRF, an inadequate immune response would favor uncontrolled cryptococcal replication and dissemination while a dysregulated immune response would control fungal burden while causing host-mediated damage to tissues and organs. AIDS patients with cryptococcal infection who are treated with antiretroviral therapy may develop a condition that is termed Immune Reconstitution Inflammatory Response (IRIS), while HIV-uninfected patients with cryptococcal meningitis may develop postinfectious inflammatory syndromes (PIIRS) following fungicidal treatment (9). As predicted by the DRF, the development of IRIS or PIIRS occurs despite control of the cryptococcal burden.

Experimental mouse models of pulmonary cryptococcal infection have revealed substantial differences in host resistance among inbred strains (10). For example, C57BL/6N inbred mice are naturally susceptible to moderately virulent strains such as *C. deneoformans* 52D (ATCC 24067) while CBA/J is naturally resistant (11). Interestingly, C57BL/6N mice develop a Th2-biased adaptive immune response characterized by eosinophilic inflammation while CBA/J mice generate a Th1-biased response characterized by expression of interferon-gamma and interleukin-12. While numerous studies have identified a contribution of individual mediators and cell types to cryptococcal immunity, the mechanisms that underlie naturally occurring differences in host resistance remain poorly understood. To investigate heritable differences in the host response to infection with moderately virulent *C. deneoformans* 52D, forward genetic analysis was performed using susceptible C57BL/6N and resistant CBA/J mice (12). This study localized natural host resistance to three intervals named *C. neoformans* susceptibility loci 1-3 (*Cnes1-3*) (12). A unique congenic mouse carrying the *Cnes2* chromosomal interval from the CBA/J inbred strain on the C57BL/6N genetic background (B6.CBA-*Cnes2*) exhibited resistance to *C. deneoformans* 52D that was characterized by a 100-fold reduction in lung fungal burden at day 35 postinfection, confirming that *Cnes2* encodes one or more host resistance genes (13).

A challenge of identifying causal genes and/or variants within a congenic chromosomal interval is to reduce the large number of potential candidate sequences (14, 15). In the case of resistant B6.CBA-*Cnes2* mice, the congenic interval spans 32.1 Mb of mouse chromosome 17 and contains 482 annotated protein-coding genes. Comparative analysis of the C57BL/6 and CBA/J genomes demonstrated that 128 of these genes have a nonsynonymous single nucleotide polymorphism (SNP) in the protein-coding region, a stop codon variation, or a splice-site variation that could regulate cryptococcal host resistance. To confine the search for causal variants, B6.CBA-*Cnes2* mice were crossed to C57BL/6N inbred mice to create sub-congenic strains, each of which encodes a subset of protein-coding sequences or regulatory elements. Here we report the creation and characterization of B6.CBA-*Cnes2b* sub-congenic mice that carry a 8.7 Mb sub-congenic segment from CBA/J on the C57BL/6N background. The *Cnes2b* sub-congenic interval encodes 175 genes of which only 61 contain a potentially relevant variant. To determine whether one or more genes within the *Cnes2b* interval regulate host resistance against *C. deneoformans* 52D infection we performed a detailed phenotypic analysis including lung fungal burden, inflammatory responses, and survival.

## Materials and Methods

### Mice

C57BL/6N mice were purchased from Inotiv and subsequently bred and maintained in the specific-pathogen-free animal facility at RI-MUHC. The B6.CBA-*Cnes2b* sub-congenic line was created by intercrossing C57BL/6N and B6.CBA-*Cnes2* mice to generate progeny that had smaller congenic fragments that were identified by genotyping by Taqman real-time PCR (ABI) with the single nucleotide polymorphism (SNP) markers rs13482444 (7.2 Mb), rs13482930 (27.0 Mb), and rs3406622636 (35.3 Mb). The limits of each sub-congenic fragment were identified by genotyping with additional SNP markers within the *Cnes2* interval. Mice with the smallest sub-congenic fragment were intercrossed to obtain a homozygous B6.CBA-*Cnes2b* line. All animals were maintained in compliance with the Canadian Council on Animal Care, and all experiments were approved by the McGill University animal care and use committee (AUP 7634).

### Intratracheal infection

*C. deneoformans* 52D (ATCC 24067) was grown and maintained on Sabouraud dextrose agar (SDA) (BD, Becton Dickinson and Company). Mice were intratracheally administered 1 × 10^4^ CFU *C. deneoformans* 52D as previously described (16) and monitored daily following surgery.

### Organ isolation and colony forming unit assay

After mice were euthanized with CO_2_, their lungs and brain were excised and placed in sterile, ice-cold phosphate-buffered saline (PBS). Tissues were then weighed, homogenized using a glass tube and pestle attached to a mechanical tissue homogenizer (Glas-Col), and plated at various dilutions on SDA. Plates were incubated at 37°C for 72 h, and CFU were counted.

### BAL fluid collection and assays

Bronchoalveolar lavage (BAL) were collected inserting a 22-gauge catheter into the trachea. A total of 4 volumes of 0.5 ml of ice-cold PBS were instilled via the catheter and subsequently aspirated. BAL was then spun (1,200 rpm; 10 min) and the supernatants were used for nitric oxide (NO) and lactate dehydrogenase (LDH) assays. NO production was assessed by the Griess assay (Sigma-Aldrich) and LDH release was quantified using the CyQUANT™ LDH Cytotoxicity Assay (Invitrogen) according to the manufacturer’s protocol.

### Blood brain barrier integrity

Mice were injected intravenously with 0.2 ml of 1% Evans Blue dye (Sigma-Aldrich) in sterile PBS. After 1 hour, the mice were sacrificed by CO_2_ inhalation, perfused with PBS, and the brains were dissected and placed in 1ml formamide for 48 hours to extract the Evans blue dye. The quantity of Evans blue dye in brain tissue was determined by spectrophotometry using a standard curve.

### Flow cytometry

For each sample a single-cell suspension was assessed by flow cytometry using an antibody panel to identify cells associated with the innate or adaptive immune response, as previously described (16).

### Total lung cytokine and chemokine production

Mice were euthanized and lungs flushed with 10 ml of ice-cold PBS. Whole lungs were homogenized in 2 ml PBS with Halt protease and phosphatase inhibitor cocktail (Thermo Scientific) using a sterilized glass tube and pestle attached to a mechanical tissue homogenizer (Glas-Col) and spun at 12,000 rpm for 20 min. Supernatants were collected, and aliquots were stored at −80^◦^C for further analysis. Total protein concentration of each sample was measured using Pierce BCA Protein Assay kit (Thermo Scientific). Levels of Cxcl1, Cxcl10, Ccl2, Ccl3, Il-1β, Il-4, Il-5, Il-6, Il-13, Il-17A, Ifn-γ and Tnf-α were quantified by multiplex ELISA using the MILLIPLEX Mouse Cytokine/Chemokine Magnetic Bead Panel (MCYTOMAG-70K, MilliporeSigma) according to the manufacturer’s protocol. Multiplex ELISA data were acquired with Luminex MAGPIX instrument and xPONENT platform, and analysis was performed on MILLIPLEX Analyst software (version 5).

### Statistical analysis

For all experiments, the mean and standard error of the mean (SEM) are shown unless otherwise stated. To test the significance of single comparisons, an unpaired Mann–Whitney test was applied with a threshold p value of 0.05. A Log-rank (Mantel-Cox) test was performed to analyze the survival curves. The significance of the difference in brain dissemination rates was determined using a Fisher’s exact test. All statistical analyses were performed with GraphPad Prism software version 10.1 (GraphPad Software Inc.).

## Results

### B6.CBA-*Cnes2b* mice display a reduced fungal burden in the lung following *C. deneoformans* 52D infection

Previous studies showed that the 32.1 Mb *Cnes2* congenic segment from CBA/J mice conferred a significant reduction in lung fungal burden following infection with *C. deneoformans* 52D compared to inbred C57BL/6N mice (13). To evaluate the influence of the smaller 8.7 Mb *Cnes2b* sub-congenic segment on *C. deneoformans* 52D infection, we first analyzed the pulmonary fungal burden in sub-congenic B6.CBA-*Cnes2b,* and inbred C57BL/6 mice at day 28 after intratracheal infection. At day 28 postinfection, the lung fungal burden of B6.CBA-*Cnes2b* was 17.4-fold lower than C57BL/6N mice (log_10_ CFU, 5.80 ± 0.19 versus 7.04 ± 0.06; P<0.0001) (Fig. 1B) showing that the *Cnes2b* sub-congenic interval localized to the distal 8.7 Mb of the *Cnes2* congenic interval is sufficient to regulate cryptococcal growth in mouse lungs.

**Figure 1.**
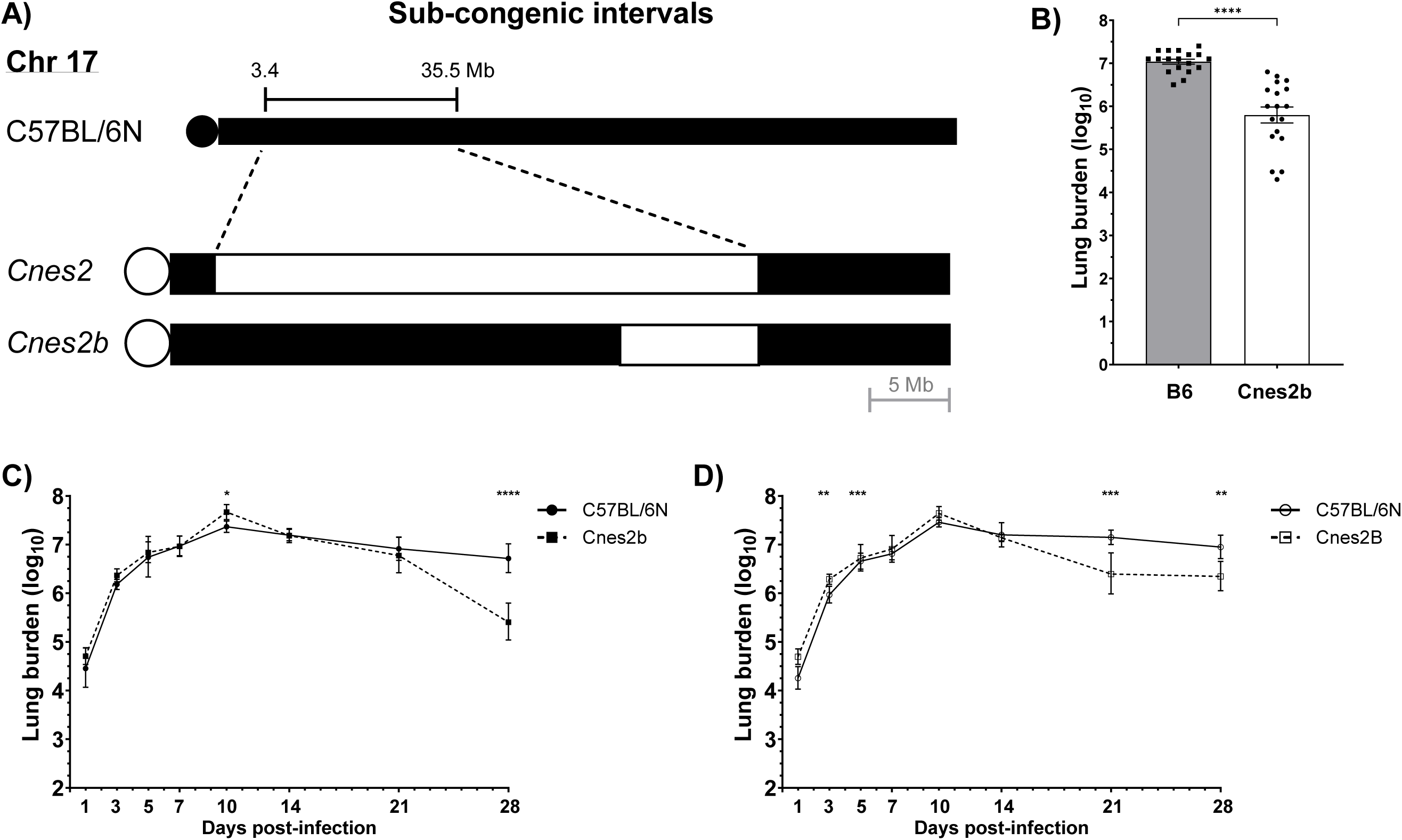
The *Cnes2b* sub-congenic interval regulates host resistance to C*. deneoformans 5*2D infection. **A)** Schematic representation of the *Cnes2b* interval on chromosome 17 (Chr 17). **B)** Lung fungal burden at 28 days post-infection with 10^4^ CFU of *C. deneoformans* 52D. Relative to inbred C57BL/6N (B6) and B6.CBA-*Cnes2b* sub-congenic mice have a significant reduction in lung fungal burden. Lung fungal burden in **C)** males and **D)** females at different days post-infection with 10 ^4^ CFU of *C. deneoformans* 52D. Mann-Whitney statistical test, n=9-30 animals per group. \**p*< 0.05,\*\**p*< 0.01, \*\*\**p*< 0.001 and **** *p*< 0.0001

The outcome of human cryptococcal disease differs between males and females (17, 18). Accordingly, to determine whether *Cnes2b* differentially regulates host resistance to experimental *C. deneoformans* infection according to sex, a comparative analysis of lung fungal burden was performed between male and female C57BL/6N and B6.CBA-*Cnes2b* mice from day 1 to day 28 postinfection. In male mice, no differences between strains were observed until day 10 postinfection when B6.CBA-*Cnes2b* had a small yet statistically significant increase in lung fungal burden compared to C57BL/6N (Fig. 1C). Subsequently, B6.CBA-*Cnes2b* male mice had a progressive reduction in lung fungal burden compared to C57BL/6N that was statistically significant at day 28 postinfection. Female B6.CBA-*Cnes2b* mice had a small yet statistically significant increase in lung fungal burden compared to C57BL/6N at day 3 and day 5 postinfection that was followed by a significant reduction at day 21 and day 28 postinfection (Fig. 1D). Comparison of male and female B6.CBA-*Cnes2b* mice showed that males had a significantly lower lung fungal burden compared to females at day 28 postinfection (Fig. S1). Taken together, these data show that *Cnes2b* regulates cryptococcal host resistance differently according to the sex of the host. Specifically, while the onset of fungal control was slightly delayed in males compared to females, the lung fungal load at day 28 postinfection was significantly lower. Thus, while *Cnes2b* sub-congenic mice have greater protection against *C. deneoformans* lung infection compared to C57BL/6N, the effect is significantly greater in males compared to females.

### B6.CBA-*Cnes2b* mice have increased mortality compared to C57BL/6 inbred mice following *C. deneoformans* 52D infection

In a previous report both C57BL/6N and B6.CBA-*Cnes2* mice with a 32.1 Mb congenic segment from CBA/J mice survived *C. deneoformans* 52D infection for 35 days (13). In the same study, both male and female B6.CBA-*Cnes2* congenic mice had a reduced *C. deneoformans* 52D burden in the lungs and spleen and a trend towards a lower rate of brain dissemination at day 35 postinfection compared to C57BL/6N inbred mice (13). In contrast to these earlier observations, characterization of *C. deneoformans* 52D infection in this study revealed an unexpected mortality rate in B6.CBA-*Cnes2b* mice. Specifically, B6.CBA-*Cnes2b* male mice began to die at day 18 postinfection and 35% had died by day 25 post-infection, while B6.CBA-*Cnes2b* female mice began to die at day 21 postinfection and 9% had died by day 25 postinfection (Fig. 2A). The mortality rate of male B6.CBA-*Cnes2b* mice was significantly greater compared to female B6.CBA-*Cnes2b* mice at day 25 postinfection (Fig. 2A) and no additional deaths occurred by day 35 postinfection. In contrast to B6.CBA-*Cnes2b*, all of the male and female C57BL/6N mice were alive at day 35 postinfection.

**Figure 2.**
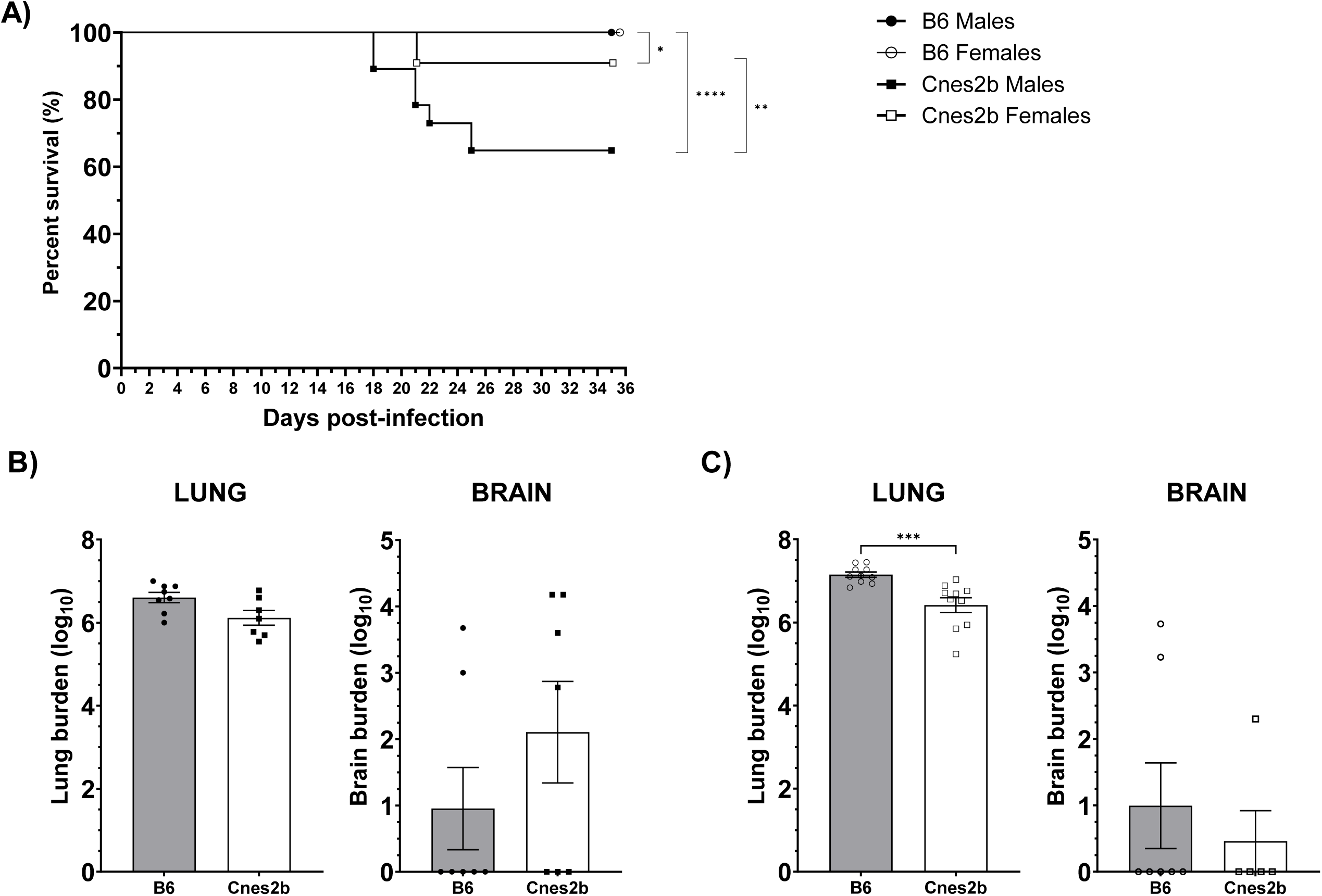
B6.CBA-*Cnes2b* sub-congenic mice have increased mortality following C. *deneoformans* 52D infection. **A)** Survival of males and females was monitored for up to 35 days after intratracheal infection with 10^4^ CFU of *C. deneoformans* 52D (n = 24-60 mice/strain, log-rank test). Fungal burden in lungs and brain prior to the onset of death for **B)** males and **C)** females at 18 or 21 days post-infection, respectively (n=9-10 mice/strain, Mann-Whitney test). * *p*< 0.05, \*\*\**p*< 0.001 and **** *p*< 0.0001.

### The increased mortality of B6.CBA-*Cnes2b* mice following *C. deneoformans* 52D infection is not associated with impaired host resistance in the lungs or brain

To determine whether the mortality of B6.CBA-*Cnes2b* mice following pulmonary *C. deneoformans* 52D infection was associated with altered host resistance to infection, lung fungal burden was quantified prior to the onset of death in males or females. At day 18 postinfection there was a nonsignificant trend towards reduced lung fungal burden in male B6.CBA-*Cnes2b* compared to C57BL/6N mice (log_10_ CFU, 6.11 ± 0.18 versus 6.60 ± 0.12; P=0.068) (Fig. 2B). At day 21 postinfection female B6.CBA-*Cnes2b* mice had a lower lung fungal burden compared to female C57BL/6N mice (log_10_ CFU, 6.42 ± 0.18 versus 7.15 ± 0.06; P<0.0001) (Fig. 2C). Taken together these data suggest that mortality of B6.CBA-*Cnes2b* mice was not due to impaired control of lung fungal burden. As meningoencephalitis is a serious and potentially fatal consequence of pulmonary cryptococcal disease, we also examined whether the *Cnes2b* sub-congenic segment derived from CBA/J caused an altered rate of fungal dissemination or growth in the brain that was associated with death. At day 18 postinfection there was no significant difference in the rate of brain dissemination between C57BL/6N and B6.CBA-*Cnes2b* males (4/7 versus 2/7; P=0.6) (Fig. 2B) or females (1/5 versus 2/7; P>0.99) (Fig. 2C). In addition, there was no significant difference in brain fungal burden between C57BL/6N and B6.CBA-*Cnes2b* males (log_10_ CFU, 0.95 ± 0.62 versus 2.10 ± 0.76; P=0.3) or females (log_10_ CFU, 0.99 ± 0.64 versus 0.46 ± 0.46; P=0.7). These data demonstrate that death of B6.CBA-*Cnes2b* mice was not associated with a greater incidence or severity of fungal meningoencephalitis.

### B6.CBA-*Cnes2b* mice have increased injury in the lungs and brain following *C. deneoformans* 52D infection

Lung injury due to infection is a complex process that is mediated by inflammation and results in alteration of the alveolar–capillary barrier (19). Nitric oxide (NO) is a potent inflammatory mediator that is produced by Nitric Oxide Synthase 2 (NOS2), an inducible enzyme in alveolar macrophages and lung epithelial cells (20). To characterize lung injury due to *C. deneoformans* 52D infection, by C57BL/6N and B6.CBA-*Cnes2b* mice, NO production was analyzed in bronchoalveolar lavage (BAL) at day 21 post-infection. Significantly higher NO production was observed in the BAL of male and female B6.CBA-*Cnes2b* compared to C57BL/6N mice (Fig. 3A). To assess changes in airway and alveolar epithelium permeability following *C. deneoformans* 52D infection, lactate dehydrogenase (LDH) was measured in BAL at day 21 postinfection. Compared to C57BL/6N, the quantity of BAL LDH was significantly greater among male and female B6.CBA-*Cnes2b* mice (Fig. 3B). Interestingly, female C57BL/6N and B6.CBA-*Cnes2b* mice also had a higher level of BAL LDH compared to male C57BL/6N and B6.CBA-*Cnes2b* mice, respectively at day 21 postinfection (Fig. 3B).

**Figure 3.**
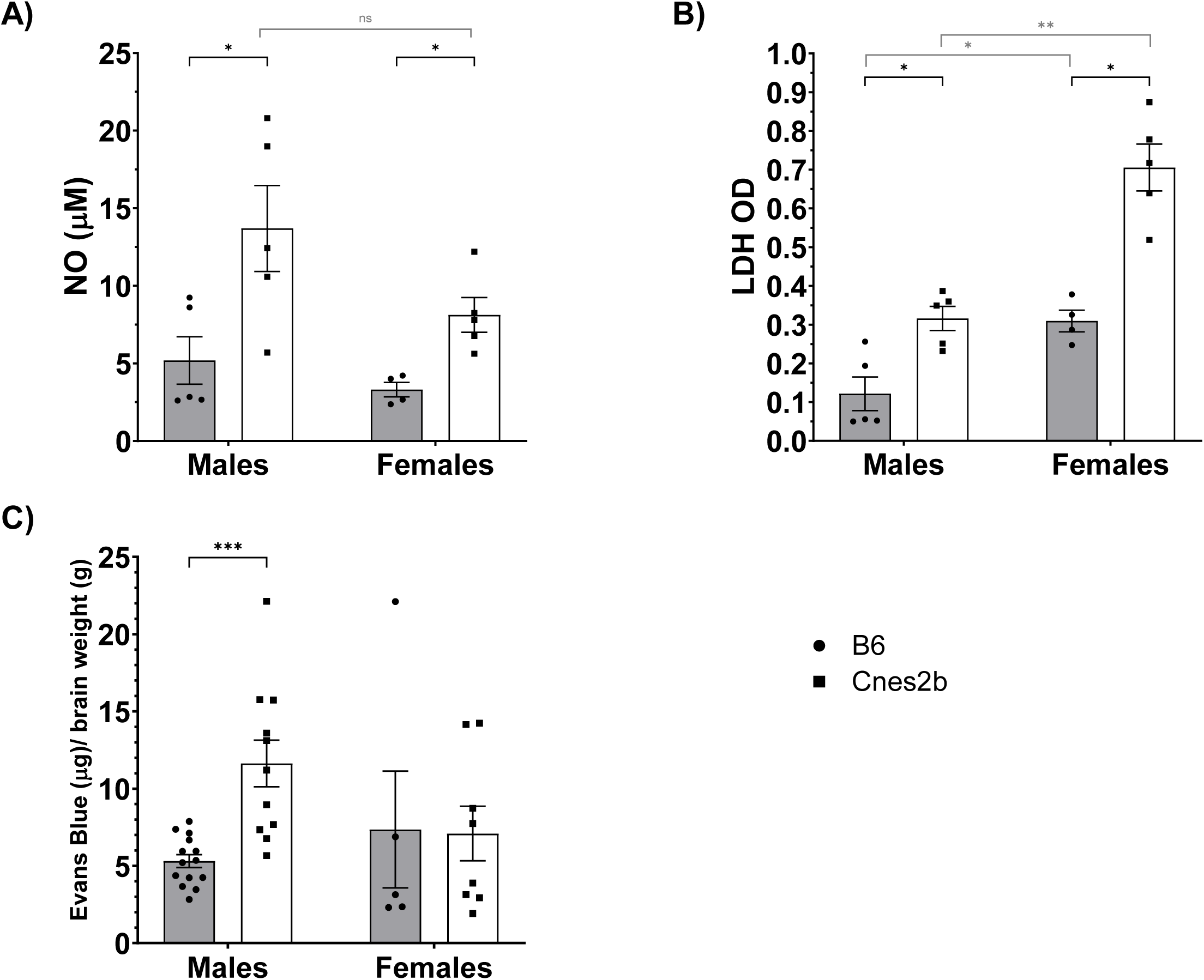
B6.CBA-*Cnes2b* subcongenic mice have increased inflammatory damage of the lungs and brain following *C. deneoformans* 52D infection. **A)** Nitric oxide (NO) and **B)** Lactate dehydrogenase (LDH) levels in bronchoalveolar lavage (BAL) at 21 days post-infection with 10^4^ CFU of *C. deneoformans* 52D. **C)** Blood-brain barrier integrity at 18 days post-infection with 10^4^ CFU of *C. deneoformans* 52D. Mann-Whitney statistical test. \**p*< 0.05, \*\**p*< 0.01, \*\*\**p*< 0.001.

Fungal dissemination to the brain from the bloodstream through various mechanisms leads to the development of cryptococcal meningitis (3). To characterize changes in blood brain barrier integrity in response to *C. deneoformans* 52D infection, Evans blue dye was quantified in the brain tissue following intravenous injection. At day 18 postinfection, male B6.CBA-*Cnes2b* sub-congenic mice had a higher quantity of Evans blue dye compared to male C57BL/6N mice (Fig. 3C). In contrast, no significant difference in the quantity of Evans blue dye was observed in the brains of female B6.CBA-*Cnes2b* and female C57BL/6N mice. These data suggest that the increased mortality of male compared to female B6.CBA-*Cnes2b* mice following *C. deneoformans* 52D infection is a consequence of increased blood brain barrier permeability.

### *Cnes2b* regulates inflammatory, Th1-, Th2-, Th17-associated cytokine and chemokine production

Previous studies have shown that *Cnes2* congenic mice that are resistant to progressive cryptococcal infection mount a Th1 pattern of adaptive immunity that is crucial for fungal clearance (13). Accordingly, to determine whether *Cnes2b* regulates host resistance to *C. deneoformans* infection through differential immune polarization, we quantified the expression of proinflammatory mediators (I1-1β, Il-6, Tnf-α) as well as Th1-type (Ifn-γ), Th2-type (Il-4, Il-5, and Il-13), Th17-type (Il-17A) cytokines and chemokines (Ccl2, Ccl3, Cxc1, Cxcl10) in total lung homogenates of B6.CBA-*Cnes2b* and C57BL/6N mice prior to infection and at day 7, 14 and 21 postinfection (Fig. 4 and 5). Prior to intratracheal *C. deneoformans* 52D challenge, all mediators were produced at a level near or below the limit of detection and no significant sex or strain differences were identified. To correlate the observed sex differences in host resistance and inflammation following *C. deneoformans* 52D infection with cytokine and chemokine production, male or female B6.CBA-*Cnes2b* and C57BL/6N mice were analyzed separately.

**Figure 4.**
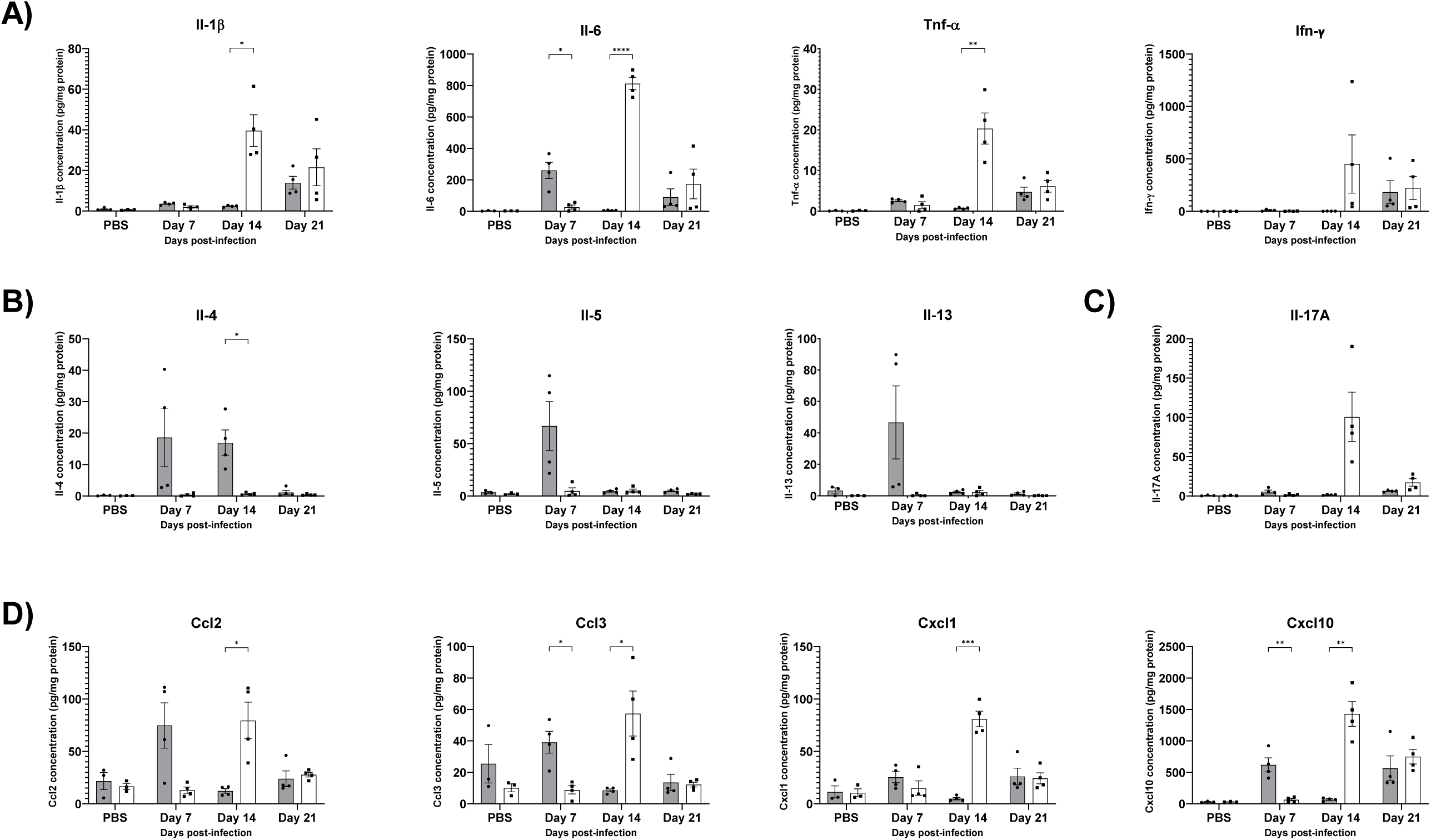
The *Cnes2b* subcongenic interval regulates lung inflammatory cytokine and chemokine production in male mice following *C. deneoformans* 52D infection. Whole lung proteins were collected at 7, 14 and 21 days post infection with 10^4^ CFU of *C. deneoformans* strain 52D from B6 (grey bars) and B6.CBA-*Cnes2b* (white bars) male mice and the quantity of **A)** pro-inflammatory immune mediators, **B)** Type 2 cytokines, **C)** Type 3 cytokines and **D)** chemokines were analyzed. Data is shown as mean ± SEM. \**p* ≤ 0.05, \*\**p*≤ 0.01, \*\*\**p* ≤ 0.001, and \*\*\*\**p* ≤ 0.0001.

**Figure 5.**
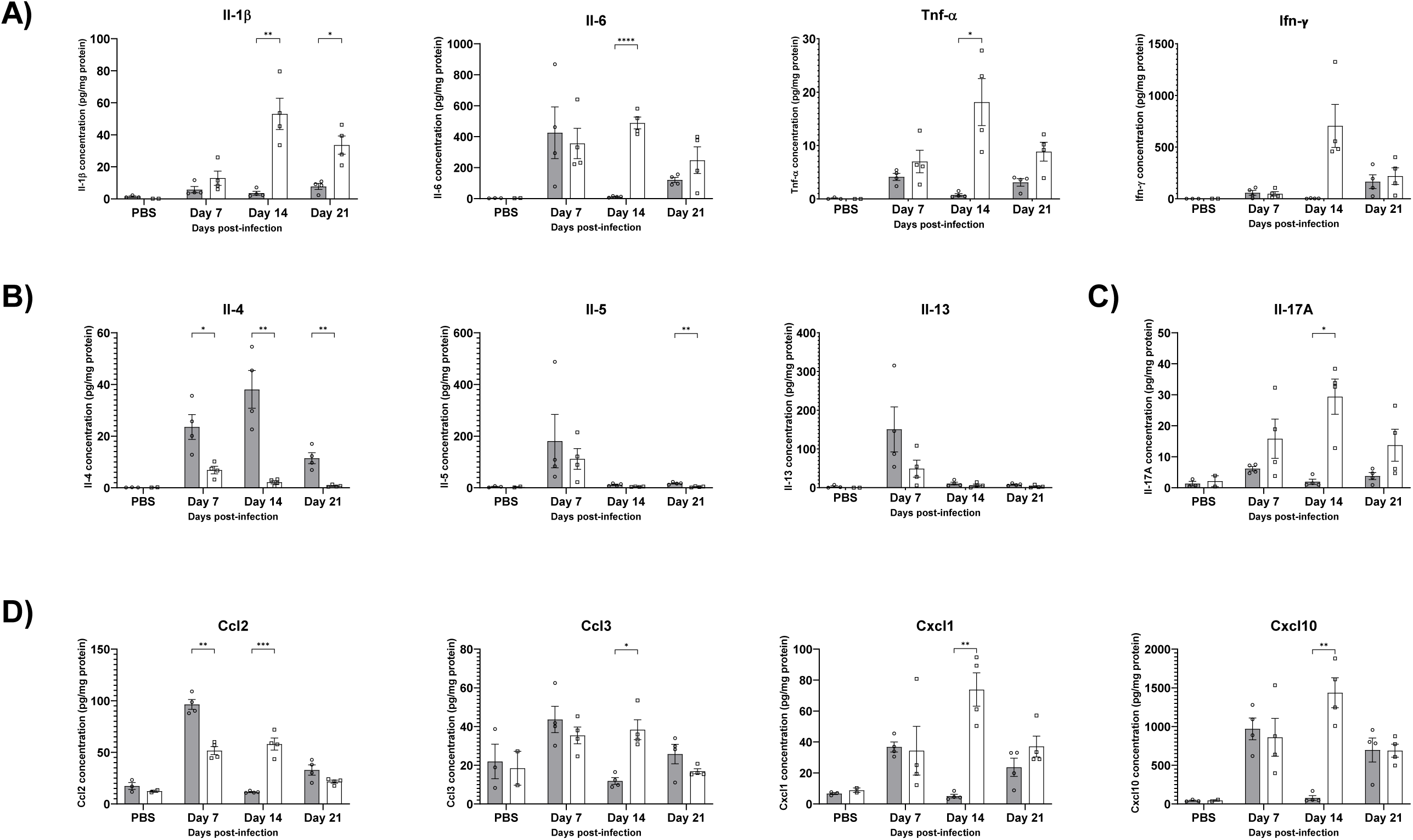
The *Cnes2b* subcongenic interval regulates lung inflammatory cytokine and chemokine production in female mice following *C. deneoformans* 52D infection. Whole lung proteins were collected at 7, 14 and 21 days post infection with 10^4^ CFU of *C. deneoformans* strain 52D from B6 (grey bars) and B6.CBA-*Cnes2b* (white bars) female mice and the quantity of **A)** pro-inflammatory immune mediators, **B**) Type 2 cytokines, **C)** Type 3 cytokines and **D)** chemokines were analyzed. Data is shown as mean ± SEM. \**p* ≤ 0.05, \*\**p*≤ 0.01, \*\*\**p* ≤ 0.001, and \*\*\*\**p* ≤ 0.0001.

Among males, the production of the proinflammatory mediators Il-1β, Il-6, and Tnf-α was significantly higher in the lungs of B6.CBA-*Cnes2b* compared to C57BL/6N mice at day 14 postinfection while Il-6 production was greater in C57BL/6N mice compared to B6.CBA-*Cnes2b* at day 7 postinfection (Fig. 4A). The production of the Th1-type cytokine Ifn-γ was at or below the detection limit at day 7 postinfection with no significant differences between B6.CBA-*Cnes2b* and C57BL/6N mice at day 14 or day 21 postinfection. A non-significant trend towards greater production of the Th2-associated cytokines Il-4, Il-5 and Il-13 was observed in C57BL/6N compared to B6.CBA-*Cnes2b* mice at day 7 postinfection with significantly higher Il-4 production in C57BL/6N mice at day 14 post-infection (Fig. 4B). The production of Il-17A was below or near the limit of detection at day 7 post-infection with higher production in B6.CBA-*Cnes2b* mice at day 14 post-infection (Fig. 4C). Finally, production of the Ccl3 and Cxcl10 chemokines was significantly greater in C57BL/6N compared to B6.CBA-*Cnes2b* mice at day 7 postinfection, and production of all measured chemokines (Ccl2, Ccl3, Cxcl1, Cxcl10) was significantly greater in B6.CBA-*Cnes2b* compared to C57BL/6N mice at day 14 postinfection (Fig. 4D).

Among females, the production of pro-inflammatory mediators was significantly higher in the lungs of B6.CBA-*Cnes2b* compared to C57BL/6N mice at day 14 postinfection (Fig. 5A) and Il-1β production was also greater in B6.CBA-*Cnes2b* compared to C57BL/6N mice at day 21 postinfection. Production of the Th1-type cytokine Ifn-γ was significantly greater in B6.CBA-*Cnes2b* compared to C57BL/6N mice at day 14 postinfection. Production of the Th2-associated cytokine Il-4 by C57BL/6N mice peaked at day 14 postinfection and was greater compared to B6.CBA-*Cnes2b* at day 7, 14, and 21 (Fig. 5B). C57BL/6N mice also had a non-significant trend towards increased Il-5 and Il-13 production compared to B6.CBA-*Cnes2b* at day 7 postinfection and a very low though statistically significant increase in Il-5 production at day 21 postinfection (Fig. 5B). B6.CBA-*Cnes2b* mice had increased Il-17A production compared to C57BL/6N at day 14 postinfection (Fig. 5C). Finally, production of all measured chemokines (Ccl2, Ccl3, Cxcl1, Cxcl10) was significantly greater in B6.CBA-*Cnes2b* compared to C57BL/6N mice at day 14 postinfection, and Ccl2 production was also increased at day 7 postinfection (Fig. 5D).

In summary, at day 14 postinfection with *C. deneoformans* 52D male and female B6.CBA-*Cnes2b* mice had increased production of pro-inflammatory mediators and chemokines in the lungs compared to C57BL/6N mice, while male and female C57BL/6N mice had increased Il-4 production compared to B6.CBA-*Cnes2b*. B6.CBA-*Cnes2b* female mice had increased expression of Ifn-γ and Il-17A at day 14 postinfection compared to C57BL/6N, while C57BL/6N female mice had increased Il-4 production at day 7 and day 21 compared to B6.CBA-*Cnes2b*. Differences in cytokine and chemokine production between male and female B6.CBA-*Cnes2b* mice were observed at day 7 postinfection (Fig. S2). These findings demonstrate that the *Cnes2b* sub-congenic interval regulates the production of inflammatory cytokines and chemokines following *C. deneoformans* 52D infection, as well as the balance of Th1-, Th17, and Th2-associated cytokines with B6.CBA-*Cnes2b* females having an increased Th1 and Th17 profile and a decreased Th2 profile compared to males.

### *Cnes2b* regulates cellular recruitment to the lung following *C. deneoformans* 52D infection

To characterize the effect of the *Cnes2b* sub-congenic interval on the cellular immune response during *C. deneoformans* 52D infection, flow cytometry analysis of whole-lung digests was performed on C57BL/6N and B6.CBA-*Cnes2b* mice at serial time points. The number of CD45^+^ cells in the lungs peaked at approximately day 14 postinfection and had decreased at day 28 postinfection. Lung cell subsets were analyzed separately for male (Fig. 6) or female (Fig. 7) B6.CBA-*Cnes2b* and C57BL/6N mice. Comparison of uninfected C57BL/6N and B6.CBA-*Cnes2b* mice showed relatively few lymphoid or myeloid cells in the lungs and no strain differences with the exception of a statistically significant increase in the small number CD4^+^ and CD8^+^ T cells in C57BL/6N females and a statistically significant increase in the small number of neutrophils in B6.CBA-*Cnes2b* females (Fig. 7).

**Figure 6.**
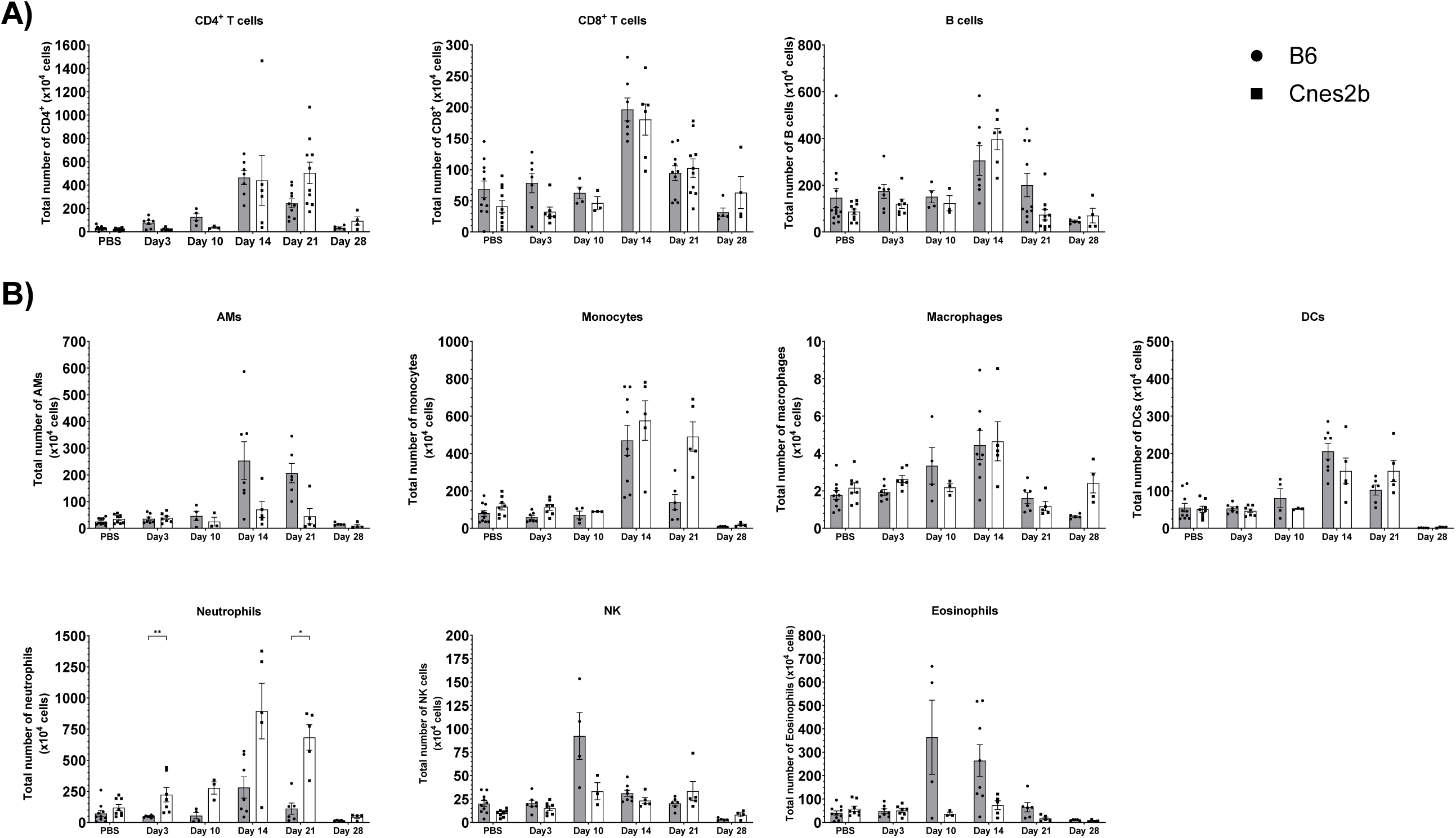
Lung flow cytometry analysis of lymphoid and myeloid cells at 14, 21 and 28 days after *C. deneoformans* 52D infection in B6 (grey bars) and B6.CBA-*Cnes2b* (white bars) male mice. **A)** Adaptive panel: CD45^+^CD3^+^CD4^+^ and CD45^+^CD3^+^CD8^+^ T lymphocytes and CD45^+^CD3^−^CD19^+^ B lymphocytes. **B)** Myeloid panel: alveolar macrophages (CD45^+^CD11c^+^F4/80^+^ SiglecF^+^), monocytes (CD45^+^SiglecF^−^ CD11b^+^Ly-6G^−^Ly-6C^hi^), macrophages (CD45^+^SiglecF^−^CD11b^+^Ly-6G^−^Ly-6C^lo-int^CD11c^−^F4/80^+^), dendritic cells (CD45^+^SiglecF^−^CD11b^+^ Ly-6G^−^Ly-6C^lo-int^CD11c^+^), neutrophils (CD45^+^SiglecF^−^CD11b^+^Ly-6G^+^Ly-6C^hi^), NK cells (CD45^+^SiglecF^−^CD11b^−^ NK1.1^+^), and eosinophils (CD45^+^CD11c^−^F4/80^int-hi^SiglecF^int-hi^). Data are shown as mean ± SEM (n = 5-7 mice/strain/time point). * *p* ≤ 0.05 and \*\**p* ≤ 0.01.

**Figure 7.**
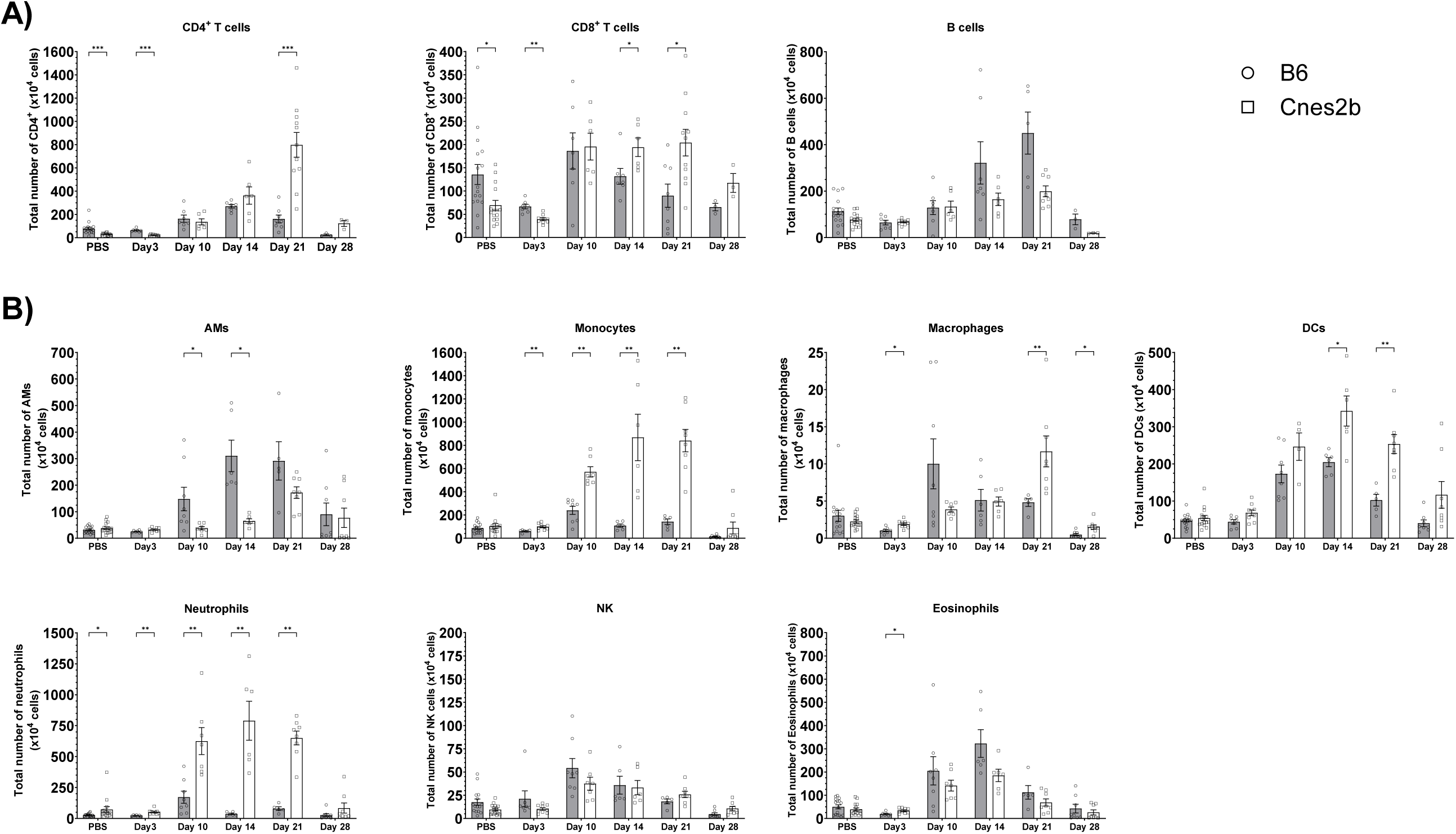
Lung flow cytometry analysis of lymphoid and myeloid cells at 14, 21 and 28 days after *C. deneoformans* 52D infection in B6 (grey bars) and B6.CBA-*Cnes2b* (white bars) female mice. **A)** Adaptive panel: CD45^+^CD3^+^CD4^+^ and CD45^+^CD3^+^CD8^+^ T lymphocytes and CD45^+^CD3^−^CD19^+^ B lymphocytes. **B)** Myeloid panel: alveolar macrophages (CD45^+^CD11c^+^F4/80^+^ SiglecF^+^), monocytes (CD45^+^SiglecF^−^ CD11b^+^Ly-6G^−^Ly-6C^hi^), macrophages (CD45^+^SiglecF^−^CD11b^+^Ly-6G^−^Ly-6C^lo-int^CD11c^−^F4/80^+^), dendritic cells (CD45^+^SiglecF^−^CD11b^+^ Ly-6G^−^Ly-6C^lo-int^CD11c^+^), neutrophils (CD45^+^SiglecF^−^CD11b^+^Ly-6G^+^Ly-6C^hi^), NK cells (CD45^+^SiglecF^−^CD11b^−^ NK1.1^+^), and eosinophils (CD45^+^CD11c^−^F4/80^int-hi^SiglecF^int-hi^). Data are shown as mean ± SEM (n = 5-7 mice/strain/time point). \**p* ≤ 0.05, \*\**p* ≤ 0.01, and \*\*\**p* ≤ 0.001.

In males, B6.CBA-*Cnes2b* mice had a significantly higher number of lung neutrophils at day 3 and 21 infection compared to C57BL/6N and had a non-statistically significant trend in the same direction at day 10 and 14. No significant differences in the number of lymphoid cells, dendritic cells, or eosinophils were observed between B6.CBA-*Cnes2b* and C57BL/6N mice following *C. deneoformans* 52D infection (Fig. 6).

In females, the total number of resident alveolar macrophages was significantly greater in C57BL/6N compared to B6.CBA-*Cnes2b* mice at day 10 and 14 postinfection, while the number of monocytes was increased in B6.CBA-*Cnes2b* mice compared to C57BL/6N at day 3, 10, 14 and 21 postinfection. B6.CBA-*Cnes2b* female mice had a significant increase in the number of lung dendritic cells at day 14 and 21 postinfection and a statistically significant increase in the number of lung macrophages at day 3, 21 and 28 postinfection. B6.CBA-*Cnes2b* female mice had a statistically significant increase in the small number of lung neutrophils and eosinophils at day 3 postinfection as well as significantly higher number of lung neutrophils at day 10, 14, and 21 postinfection compared to C57BL/6N. Interestingly, C57BL/6N females had a statistically significant increase in the small number of lung CD4^+^ and CD8^+^ T cells at day 3 postinfection while B6.CBA-*Cnes2b* mice had a significantly higher number of CD4^+^ T cells at day 21 postinfection and a significantly higher number of CD8^+^ T cells at day 21 and 28 postinfection. No significant differences in the number of B cells or NK cells was observed between B6.CBA-*Cnes2b* and C57BL/6N female mice following *C. deneoformans* 52D infection (Fig. 7).

Taken together, these findings indicate that the *Cnes2b* sub-congenic segment has a significant effect on lung recruitment of neutrophils in male mice and several myeloid cell subsets in female mice after *C. deneoformans* 52D infection. Specifically, *Cnes2b* enhances the recruitment of neutrophils, as well as inflammatory monocytes that subsequently differentiate into macrophages and dendritic cells, in females throughout the course of *C. deneoformans* 52D infection. *Cnes2b* increased the recruitment of CD4^+^ or CD8^+^ T cells in females, but had no effect on B cells or NK cells in either sex after *C. deneoformans* 52D infection.

## Discussion

Previous experimental studies established the *Cnes2* congenic interval from the naturally resistant CBA/J inbred strain regulates host resistance to progressive *C. deneoformans* 52D infection (13). *Cnes2* spans 32.1 Mb and encodes 482 annotated protein-coding genes. Comparative analysis of *Cnes2* between CBA/J and C57BL/6N revealed potentially important sequence variation in 128 of these genes. The large number of variants encoded by *Cnes2* poses a significant challenge to the identification of causal genes. To reduce the number of potential candidates that regulate host resistance, subcongenic mice were derived by outcrossing B6.CBA-*Cnes2* to C57BL/6N. The B6.CBA-*Cnes2b* sub-congenic strain carries 8.7 Mb from the telomeric region of *Cnes2* that encodes 175 annotated protein-coding genes. Comparative analysis of C57BL/6N and CBA/J showed that 61 of these genes contain either a nonsynonymous single nucleotide polymorphism (SNP) in the protein-coding region, a stop codon variation, or a splice-site SNP. Intratracheal infection with *C. deneoformans* 52D demonstrated that B6.CBA-*Cnes2b* mice have a significantly lower fungal burden at day 28 postinfection compared to C57BL/6N. Thus, despite markedly reduced genetic complexity, B6.CBA-*Cnes2b* subcongenic mice have enhanced host resistance against *C. deneoformans* 52D. This finding focuses the search for causal variants but it does not exclude the possibility that additional host resistance genes may be located proximally to the *Cnes2b* sub-interval (13). In fact, high-resolution analysis of quantitative traits in mice typically partitions single QTLs into multiple closely linked QTLs that may have additive or opposite effects (14, 21, 22).

To determine whether *Cnes2b* alters host resistance during the innate or the adaptive phase of the immune response, lung fungal burden was determined at serial time points after *C. deneoformans* 52D infection. Male and female B6.CBA-*Cnes2b* mice were analyzed separately and compared to C57BL/6N mice to identify potential sex-dependent differences that are regulated by *Cnes2b*. Somewhat unexpectedly, female B6.CBA-*Cnes2b* mice had a very small yet reproducible increase in lung fungal burden during the first 3-5 days postinfection compared to C57BL/6N, and a similar observation was made in male B6.CBA-*Cnes2b* mice at day 10 postinfection. Conversely, female and male B6.CBA-*Cnes2b* mice had a significant reduction of lung fungal burden compared to C57BL/6N beginning at day 21 and day 28 postinfection, respectively. These observations suggest that while *Cnes2b* may have a subtle effect on initial or innate control of pulmonary cryptococcal infection with a delayed onset in males, its predominant effect is to enhance subsequent control of lung fungal burden during the adaptive phase of the immune response. During the initial phase of infection, signals from the innate immune system are required for induction of adaptive immune responses. Microbial pattern recognition receptors (PRRs) and dendritic cells (DCs) are key cellular and molecular links between innate and adaptive immunity (23). Given the differences in lung fungal burden at early and late time points, it is tempting to speculate that *Cnes2b* may regulate the function of PRRs and/or DCs during *C. deneoformans* 52D infection in a way that is slightly permissive to fungal replication at early time points while facilitating more effective fungal burden control at later time points.

Observational and epidemiological studies have documented a higher incidence and poorer outcomes of human cryptococcal infection in males (17). The precise mechanisms that account for the sex difference remain unknown although experimental studies in males have suggested that the hormonal environment may affect pathogen virulence or the presence of an inherent T-cell defect may affect the host response of males (17, 24). In the current study, *Cnes2b* regulated certain aspects of the host response differently in males compared to females. First, male B6.CBA-*Cnes2b* mice had a lower lung fungal burden at day 28 postinfection compared to females. Second, male B6.CBA-*Cnes2b* mice experienced a higher mortality rate compared to females after experimental *C. deneoformans* 52D infection. Importantly, no significant differences in brain fungal burden were observed between male or female B6.CBA-*Cnes2b* and C57BL/6N mice prior to death, indicating that mortality was not attributable to increased dissemination or replication of *C. deneoformans* 52D in either sex. Evaluation of blood brain barrier integrity did show increased dye extravasation into the brains of male B6.CBA-*Cnes2b* mice, suggesting that death was associated with more severe central nervous system injury. *Cnes2b* also regulated inflammatory cell numbers in the lung differently in male and female mice. Prior to infection no significant differences were observed in the number of lung leukocytes between male B6.CBA-*Cnes2b* and C57BL/6N mice, while female B6.CBA-*Cnes2b* mice had a small decrease in the number CD4^+^ and CD8^+^ T cells and a very small increase in the number of neutrophils compared to C57BL/6N. Following *C. deneoformans* 52D infection there was no significant difference in the number of lung leukocytes between male B6.CBA-*Cnes2b* and C57BL/6N mice with the exception of increased neutrophils, while female B6.CBA-*Cnes2b* mice had significant increases in both lymphoid (CD4^+^ and CD8^+^ T cells) and myeloid (neutrophils, monocytes, macrophages, dendritic cells) cell subsets and a decreased number of alveolar macrophages compared to C57BL/6N. Taken together these data show that *Cnes2b* has a greater effect on lung cell numbers in female mice compared to male mice, both in the absence of infection as well as after intratracheal challenge with *C. deneoformans* 52D.

Certain aspects of the host response were regulated by *Cnes2b* in both males and females including increased production of inflammatory cytokines and chemokines, decreased production of Il-4, and signs of acute lung injury at day 14 postinfection relative to C57BL/6N. Il-17A production was also increased in both male and female B6.CBA-*Cnes2b* mice compared to C57BL/6N but was statistically significant only in females.

The negative consequences of an excessive host response are clearly described in the damage response framework of infectious disease pathogenesis (25). In the setting of cryptococcal infection, control of fungal burden may be complicated by an immune reconstitution inflammatory syndrome or a postinfectious inflammatory syndrome (PIIRS) in HIV-infected and HIV-uninfected patients, respectively (9). In the current study, B6.CBA-*Cnes2b* mice had more effective control of lung fungal burden that was associated with a heightened host inflammatory response, increased lung and brain damage, as well as excess mortality compared to C57BL/6N. These experimental findings closely resemble the clinical manifestations of individiuals with cryptococcal IRIS/PIIRS. B6.CBA-*Cnes2b* mice showed increased pulmonary recruitment of myeloid and lymphoid cells. Previous experimental studies have implicated a heightened inflammatory response and a pathological role for CD4^+^ T cells the pathogenesis of lethal cerebral cryptococcosis (26, 27). Thus, it is possible that identification of genes encoded by *Cnes2b* that regulate host responsiveness may uncover valuable mechanistic insights into the development of pathological host inflammation in the lungs and brain.

This study has several limitations. First, the unexpected mortality in B6.CBA-*Cnes2b* mice infected with *C. neoformans* 52D precluded a complete evaluation of the host response at all time points. As a result, the fungal burden as well as cytokine and chemokine production at day 28 may be subject to a survivor bias, particularly among male mice that had a higher mortality rate. Second, the analysis of lung and brain injury was confined to selected measures of tissue inflammation and vascular permeability and further characterization would be required to determine whether death is primarily attributable to failure of the respiratory or central nervous system. Third, additional studies will be required to determine whether enhanced lymphoid and/or myeloid cell recruitment also contributes to central nervous system injury in B6.CBA-*Cnes2b* mice.

In summary, the data in this report demonstrate pleiotropic and sex-dependent effects of the *Cnes2b* interval on the host immune response to pulmonary *C. deneoformans* 52D infection. Although the exact regulatory mechanisms remain to be defined, we hypothesize that the underlying genetic factors encoded by the *Cnes2b* interval function during the transition from innate to adaptive immunity. Definitive identification of the *Cnes2b* genes that regulate cryptococcal infection in mice may translate into novel therapeutic targets that could reduce the human disease burden.

## Acknowledgements

We thank T. Mary Fujiwara for critical review of the manuscript.

This study was supported by grants from the Canadian Institutes of Health Research (FRN 159588; STQ), the Research Institute of the McGill University Health Centre Foundation (STQ), and the J.T. Costello Memorial Bequest (STQ).

